# Nanog organizes transcription bodies

**DOI:** 10.1101/2022.06.13.495463

**Authors:** Ksenia Kuznetsova, Martino Ugolini, Edlyn Wu, Manan Lalit, Haruka Oda, Yuko Sato, Hiroshi Kimura, Florian Jug, Nadine Vastenhouw

## Abstract

The localization of transcriptional activity in specialized transcription bodies is a hallmark of gene expression in eukaryotic cells. How proteins of the transcriptional machinery come together to form such bodies, however, is unclear. Here, we take advantage of two large, isolated, and long-lived transcription bodies that reproducibly form during early zebrafish embryogenesis, to characterize the dynamics of transcription body formation. Once formed, these transcription bodies are enriched for initiating and elongating RNA polymerase II, as well as the transcription factors Nanog and Sox19b. Analyzing the events leading up to transcription, we find that Nanog and Sox19b cluster prior to transcription, and independently of RNA accumulation. The clustering of transcription factors is sequential; Nanog clusters first, and this is required for the clustering of Sox19b and the initiation of transcription. Mutant analysis revealed that both the DNA-binding domain, as well as one of the two intrinsically disordered regions of Nanog are required to organize the two bodies of transcriptional activity. Taken together, our data suggests that the clustering of transcription factors dictates the formation of transcription bodies.

**HIGHLIGHTS:**

- Transcription factors cluster prior to, and independently of transcription
- Nanog organizes transcription bodies: it is required for the clustering of Sox19b as well as RNA polymerase II
- This organizing activity requires its DNA binding domain as well as one of its intrinsically disordered regions
- Transcription elongation results in the disassembly of transcription factor clusters

---

Transcription by RNA polymerase II (RNA Pol II) takes place in concentrated areas in the nucleus, also called transcription bodies (Cisse et al. 2013; Boehning et al. 2018; Cho et al. 2018; Ghamari et al. 2013; Iborra et al. 1996; Jackson et al. 1998; Huang et al. 2021; Hadzhiev et al. 2021). Recently, it has been shown that clusters of RNA Pol II and Mediator localize to such transcription bodies (Cho et al. 2018). Transcription factors also form clusters in the nucleus (Tsai et al. 2017; Mir et al. 2017; Nair et al. 2019; Sabari et al. 2018; Li et al. 2020; Ma et al. 2021). These can recruit Mediator (Boija et al. 2018) and even RNA Pol II in artificial systems (Chong et al. 2018), but it remains unclear how transcription bodies form *in vivo*.

## RNA Pol II transcription localizes to two transcription bodies when zebrafish genome activates

In zebrafish, bulk zygotic transcription starts ∼3 hr post-fertilization, around the tenth cell division (Aanes et al. 2011; Heyn et al. 2014; Harvey et al. 2013; D A Kane and Kimmel 1993; White et al. 2017; Pauli et al. 2012) (Figure 1A). Using sensitive techniques, however, transcripts have now been detected as early as 64-cell stage (Heyn et al. 2014; White et al. 2017) (Figure 1A). To characterize the spatial organization of transcription onset in zebrafish embryos, we visualized the initiating and elongating form of RNA Pol II (RNA Pol II phosphorylated on Serine 5 (RNA Pol II Ser5P) and Serine 2 (RNA Pol II Ser2P), respectively) during zygotic genome activation. We injected 1-cell stage embryos with <u>f</u>ragments of an <u>a</u>nti<u>b</u>ody (Fab) against RNA Pol II Ser5P and Ser2P as described before (Hilbert et al. 2021; Sato et al. 2019), let them develop to the desired stage and imaged embryos on a confocal Spinning Disk microscope (Figure 1B). Following embryos from 32-cell stage onward, we first detected transcriptional activity during the 64-cell stage (Figure 1B, Movie S1). Initiating RNA Pol II was followed by the appearance of elongating RNA Pol II (Figure 1B). Analyzing the appearance of Pol II Ser5P and Ser2P in locations where they coincide confirmed this (Figure 1C). Reminiscent of RNA Pol II clusters seen in mammalian cells (Iborra et al. 1996; Jackson et al. 1998; Boehning et al. 2018; Cho et al. 2018), RNA Pol II activity localized to distinct transcription bodies in the nucleus. In contrast to previous studies in which many transcription bodies were observed, however, productive transcription in the zebrafish embryo is initially limited to two transcription bodies. We continued to observe these two prominent transcription bodies in subsequent stages (128- and 256-cell stage are shown in Figure 1B,C; also see Movies S2,3), and here too, the elongating form of RNA Pol II follows the initiating form. Transcription bodies appear earlier in the cell cycle in consecutive stages (Figure 1B-D). They grow during the cell cycle (Figure 1E) and grow larger during later stages (Figure 1F). We note that from the ninth cell cycle onward, more than two clusters of initiating RNA Pol II can be observed (Figure 1B). Remarkably, these additional initiating RNA Pol II clusters do not accumulate any elongating RNA Pol II (Figure 1B,G), suggesting that transcription elongation initially takes place only in two transcription bodies. We conclude that during zebrafish genome activation, productive transcription starts in two transcription bodies in the nucleus. These transcription bodies are isolated, large, long-lived, and appear at a predictable time during development.

**Figure 1.**
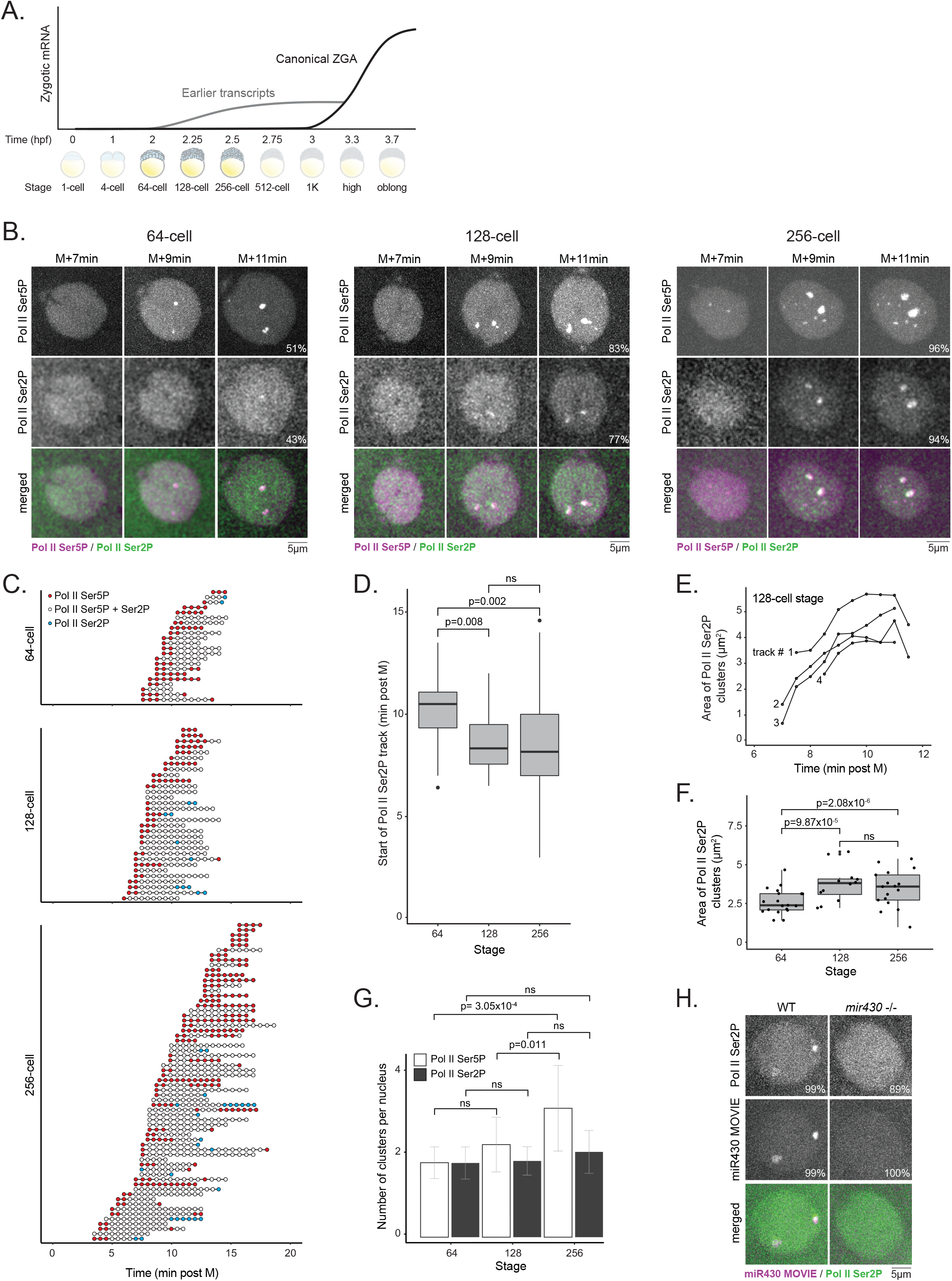
RNA Pol II transcription localizes to two transcription bodies when zebrafish genome activates. **A**. Schematic representation of zebrafish zygotic genome activation. hpf, hours post fertilization. **B**. Visualization of initiating and elongating RNA polymerase II (Pol II Ser5P and Pol II Ser2P, respectively) with fragmented antibodies (Fabs). Representative images of individual nuclei from 64, 128 and 256-cell stage embryos, extracted from spinning disk confocal microscopy time lapse at 7, 9 and 11 minutes after mitosis. The percentage of nuclei with at least one Pol II (Ser5P or Ser2P) cluster is indicated. **C**. Tracks of transcription bodies at 64-, 128-, and 256-cell stage. The presence of Pol II Ser5P, Ser2P, or both, is indicated by red, blue, and white circles, respectively. Time on the x-axis in minutes after mitosis. **D**. Boxplot representing the time of start of Ser2P track in minutes after mitosis at 64-, 128 and 256-cell stage. Filled circles represent outliers. **E**. Evolution of Pol II Ser2P area (in μm^2^) for the four earliest detected tracks of Pol II Ser2P clusters at 128-cell stage. Time on the x-axis in minutes after mitosis. **F**. Area of Pol II Ser2P clusters at the midpoint of the indicated interphase in μm^2^. **G**. Average number of Pol II Ser5P and Pol II Ser2P clusters in active nuclei (defined by the presence of at least one Pol II cluster). Mean ± SD. For B-G, 64-cell stage (N=6, n=35), 128-cell stage (N=6, n=69), 256-cell stage (N=6, n=97). **H**. Visualization of Pol II Ser2P and miR430 MOVIE signal in wildtype and *mir430* mutant embryos. Shown are representative images of nuclei at the midpoint of interphase at 128-cell stage. The percentage of nuclei with at least one Pol II Ser2P cluster is indicated. For wt, N=3, n=111; for *mir430* mutant N=3, n=72. Pairwise non-parametric Wilcoxon tests were performed, ns indicates P > 0.05. All images represent maximum intensity projections (MIPs) through the Z-stack.

Among the most expressed genes in early zebrafish embryos is *mir430* (Heyn et al. 2014; White et al. 2017), which is encoded by a cluster of *mir430* genes. We and others have previously shown that the first two transcription bodies that characterize the onset of transcription in zebrafish embryos localize to the *mir430* locus (Hilbert et al. 2021; Hadzhiev et al. 2019; Chan et al. 2019; Sato et al. 2019). To confirm this, we visualized the two transcription bodies using the RNA Pol II Ser2P Fab, and miR430 transcripts using <u>Mo</u>rpholino <u>Vi</u>sualization of <u>E</u>xpression (MOVIE; Hadzhiev et al. 2019) at 128-cell stage. As expected, RNA Pol II Ser2P signal and miR430 transcripts coincide (Figure 1H). Next, to determine whether the *mir430* locus is required for the formation of the first transcription bodies, we generated a mutant in which we deleted the complete *mir430* locus (Figure S1). In the *mir430* deletion mutant, both RNA Pol II Ser2P and miR430 MOVIE signals were absent in most nuclei (Figure 1H). This shows that the two transcription bodies that characterize the beginning of transcription in zebrafish embryos are seeded by the *mir430* locus.

## Transcription factors cluster prior to, and independent of transcription

Transcription bodies have been observed in multiple systems (Cisse et al. 2013; Boehning et al. 2018; Cho et al. 2018; Ghamari et al. 2013; Iborra et al. 1996; Jackson et al. 1998; Huang et al. 2021; Hadzhiev et al. 2021). It is unclear, however, what organizes their formation. Transcription factors have also been shown to form clusters in the nucleus (Tsai et al. 2017; Mir et al. 2017; Nair et al. 2019; Sabari et al. 2018; Li et al. 2020; Ma et al. 2021) and because transcription factors are well-known to recruit transcriptional machinery (Kadonaga et al. 1988; Ptashne and Gann 1997; Lu, Portz, and Gilmour 2019; Nair et al. 2019; Boija et al. 2018; Chong et al. 2018; Kwon et al. 2013), it seems likely that transcription factor clusters organize bodies of transcriptional activity. To investigate this, we visualized Nanog, Sox19b and Pou5f3 which have previously been shown to be required for the transcription of *mir430* (Lee et al. 2013; Leichsenring et al. 2013; Onichtchouk et al. 2010). We injected RNA encoding each of these transcription factors fused to mNeonGreen (mNG) at the 1-cell stage and visualized each transcription factor in combination with the initiating form of RNA Pol II (Figure 2A, and Movies S4-6). We focused our analysis on 128-cell stage embryos because at this stage most nuclei show at least one (Figure 1B), and on average two (Figure 1G) transcription bodies, which we know to be seeded by the *mir430* locus (Figure 1H). We found that Nanog forms multiple clusters in the nucleus (Figure 2A, Nanog; Movie S4). Exactly two of these coincide with the two *mir430* transcription bodies. Sox19b formed only two clusters in the nucleus (Figure 2A, Sox19b; Movie S5). These both coincide with the two transcription bodies. Pou5f3, like Nanog, formed multiple clusters in the nucleus (Figure 2A, Pou5f3; Movie S6). However, unlike the Nanog clusters, these did not coincide with the two transcription bodies. While this does not mean that Pou5f3 is not present in these transcription bodies, it may reflect the minor role of Pou5f3 in *mir430* transcription (Lee et al. 2013). We conclude that the two *mir430* transcription bodies are enriched for Nanog and Sox19b, but not for Pou5f3.

**Figure 2.**
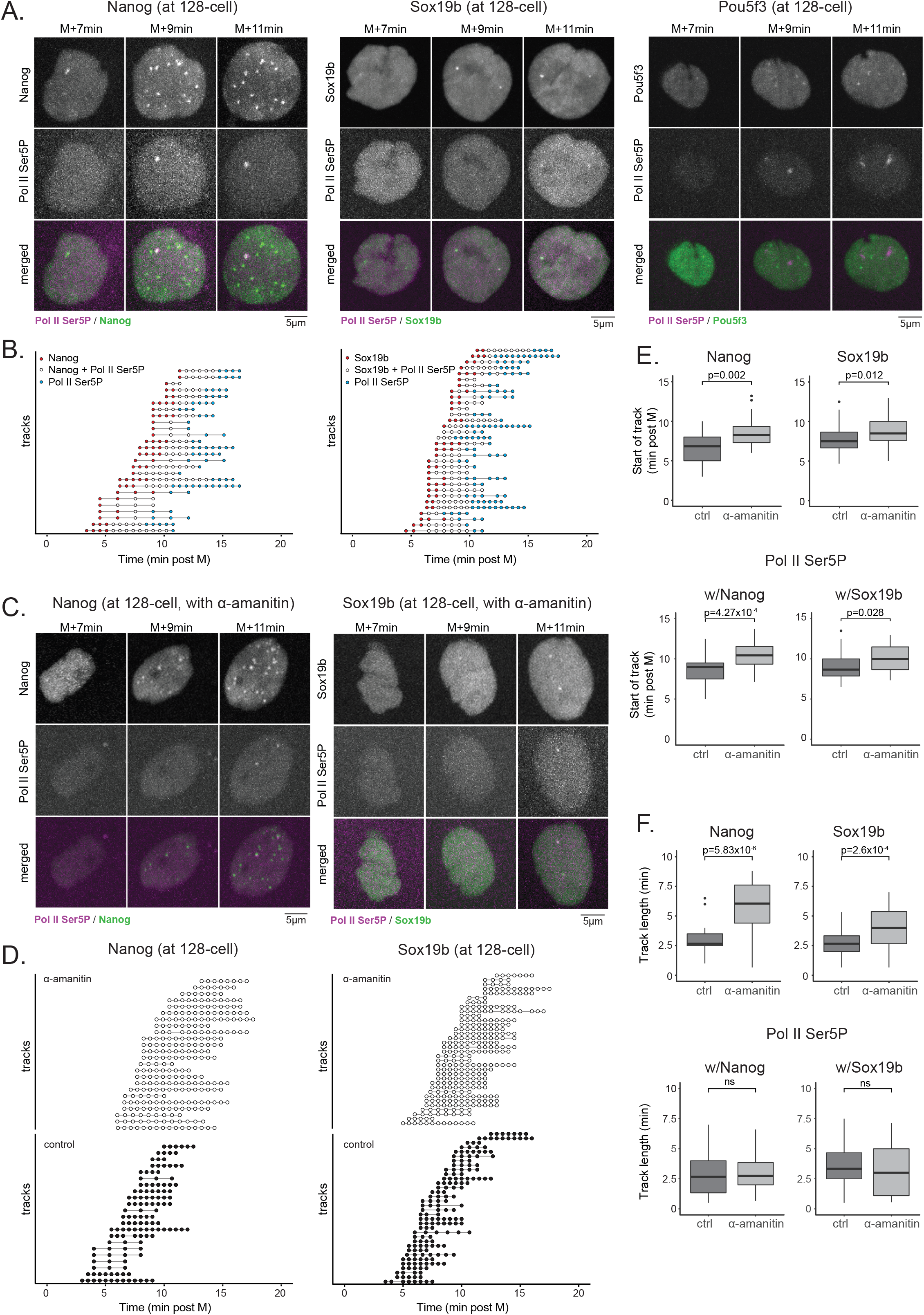
Transcription factors cluster prior to, and independent of transcription. **A**. Visualization of Nanog, Sox19b, and Pou5f3, in combination with the initiating form of RNA Pol II. Representative images of individual nuclei from 128-cell stage embryos are shown, extracted from spinning disk confocal microscopy time lapse at 7, 9 and 11 minutes after mitosis. **B**. Tracks of transcription factor and Pol II Ser5P clusters at 128-cell stage. The presence of TF (Nanog (left panel), or Sox19b (right panel)), TF and Pol II Ser5P, or Pol II Ser5P alone is indicated by red, white, and blue circles, respectively. Time on the x-axis in minutes after mitosis. For A and B, Nanog (N=4, n=24), Sox19b (N=4, n=37), Pou5f3 (N=3, n=20). **C**. Visualization of transcription factors (Nanog (left panel), Sox19b (right panel)), and Pol II Ser5P at 128-cell stage in a-amanitin treated embryos. Representative images are shown, extracted from spinning disk confocal microscopy time lapse at 7, 9 and 11 minutes after mitosis. **D**. Tracks of Nanog (left panel), and Sox19b (right panel)) at 128-cell stage in a-amanitin treated (white) and control (black) embryos. Time on the x-axis in minutes after mitosis. For C and D, Nanog (N=3, n=27), Sox19b (N=4, n=38). **E**,**F**. The effect of transcription inhibition (by a-amanitin) on start (E) and length (F) of Nanog, Sox19b, and Pol II Ser5P tracks (in minutes post mitosis). Pol II Ser5P data is shown separately for nuclei in which Nanog and Sox19b were visualized. Pairwise non-parametric Wilcoxon tests were performed, ns indicates P > 0.05. All images represent maximum intensity projections (MIPs) through the Z-stack.

To determine the relationship between the clustering of transcription factors and transcription, we analyzed the appearance of Nanog and Sox19b signal in relation to the appearance of RNA Pol II Ser5P signal in the *mir430* transcription bodies (Figure 2B). This showed that both Nanog and Sox19b cluster before RNA Pol II Ser5P. Thus, transcription factors can cluster independently of transcription initiation. This is supported by the observation that most Nanog clusters we observe do not coincide with RNA Pol II Ser5P (Figure 2A). To further test the role of transcriptional activity in transcription factor clustering, we inhibited transcription elongation by injecting alpha-amanitin in 1-cell stage embryos. Amanitin-injected embryos arrested at sphere stage as previously described (Pálfy et al. 2020; Lee et al. 2013; Joseph et al. 2017), and no RNA Pol II Ser2P signal was observed (data not shown, and (Sato et al. 2019)). Nanog and Sox19b, however, were still able to cluster (Figure 2C-E and Movies S7,8). We conclude that transcription factors cluster prior to, and independent of transcription elongation.

## RNA accumulation results in dissociation of transcription factor clusters

The role of RNA in cluster formation is multi-faceted. While RNA is known to facilitate cluster formation (Falahati et al. 2016; Quinodoz et al. 2021; Banerjee et al. 2017), higher concentrations of RNA have been associated with the dissolution of clusters (Maharana et al. 2018; Henninger et al. 2021; Shao et al. 2021; Nozawa et al. 2017). To investigate the effect of RNA accumulation on Nanog and Sox19b clusters, we compared the time after anaphase at which we started to see transcription factor clusters in amanitin- and control-injected embryos (Figure 2D,E). For both Nanog and Sox19b, we found that in the absence of transcription, signal appeared significantly later. In line with the observation that RNA Pol II Ser5P clustering follows transcription factor clustering, the appearance of RNA Pol II Ser5P (initiation) clusters was also delayed in the absence of transcription elongation (Figure 2E). Thus, our data suggests that RNA accumulation is not required for transcription factor clustering, but it does facilitate the process. Next, we investigated the impact of RNA accumulation on the dissolution of transcription factor clusters by plotting the lifetime of Nanog, Sox19b and RNA Pol II Ser5P clusters in amanitin- and control-injected embryos (Figure 2F). For both Nanog and Sox19b, cluster lifetime was significantly increased in the absence of transcription. The lifetime of RNA Pol II Ser5P clusters was not affected, which we assume is a consequence of the complex relationship between transcription elongation and initiation. We conclude that RNA facilitates the formation of transcription factor clusters, while accumulation of RNA causes them to dissolve. This is in agreement with the previously described role of RNA in cluster formation and dissolution.

## Nanog organizes transcription bodies

The observation that both Nanog and Sox19b cluster prior to and independent of transcription elongation, raises the question in what order the two transcription factors cluster. To investigate this, we plotted the appearance of Nanog, Sox19b, and RNA Pol II Ser5P in the *mir430* transcription bodies relative to the anaphase of each cell cycle (Figure 3A). This revealed that Nanog clusters first, followed by Sox19b and RNA Pol II Ser5P. Averaging and comparing the time at which Nanog, Sox19b and RNA Pol II Ser5P can be observed confirmed this (Figure 3B). Because Nanog clusters first, we then asked whether Nanog is required for the clustering of Sox19b. For this, we compared Sox19b clustering in wildtype and *Nanog* mutant embryos using a *Nanog* mutant that we previously characterized (Veil et al. 2018; Pálfy et al. 2020). As before, we visualized Sox19b by injecting RNA encoding Sox19b fused to mNeongreen into 1-cell stage embryos. In the presence of Nanog (wildtype embryos), we consistently observed two clusters of Sox19b. In the absence of Nanog (*Nanog* mutant), however, we did not observe such clusters (Figure 3C, Movies S9,10). This suggests that Nanog is required for the clustering of Sox19b. Next, to determine whether in absence of Nanog RNA Pol II Ser5P could still be detected, we injected Fabs into embryos of wildtype and *Nanog* mutant embryos. This clearly showed that in the absence of Nanog, no RNA Pol II Ser5P signal was detected (Figure 3D, Movies S11,12). We conclude that Nanog is required for the successive clustering of Sox19b and RNA Pol II Ser5P, and thus for the organization of transcription bodies.

**Figure 3.**
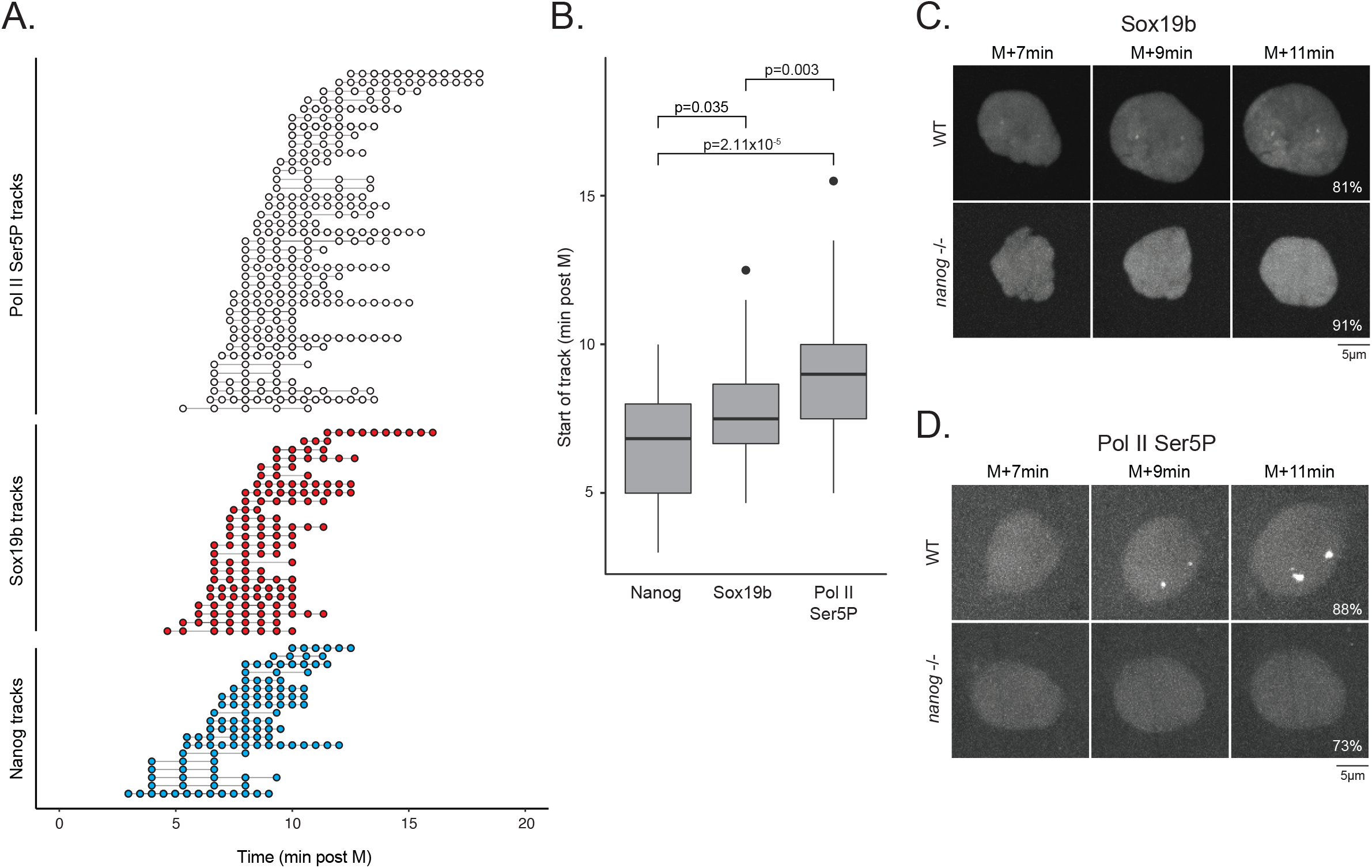
Nanog organizes transcription bodies. **A**. Tracks of Nanog (blue), Sox19b (red), and Pol II Ser5P (white), extracted from 128-cell stage embryos as represented in Figure 2A. **B**. Average start time (in minutes after Mitosis) of Nanog, Sox19b and Pol II Ser5P tracks shown in A. **C**. Visualization of Sox19b in wt (upper panel, N=4, n=38) and Nanog mutant (lower panel, N=3, n=30) embryos. **D**. Visualization of Pol II Ser5P in wt (upper panel, N=3, n=41) and Nanog mutant (lower panel, N=4, n=58) embryos. Representative images are shown, extracted from spinning disk confocal microscopy time lapse at 7, 9 and 11 minutes after mitosis. All images represent maximum intensity projections (MIPs) through the Z-stack. In C-D, the percentage of nuclei with the indicated pattern is indicated. Pairwise non-parametric Wilcoxon tests were performed, ns indicates P > 0.05.

## Nanog DBD as well as IDR are required to organize transcription bodies

The essential role of Nanog in the formation of the two *mir430* transcription bodies provided us with the opportunity to dissect which Nanog domains are required for the formation of transcription bodies. Nanog is a homeodomain-containing transcription factor, mostly known for its role in maintaining pluripotency (Silva et al. 2009; Chambers et al. 2003; Boyer et al. 2005; Mitsui et al. 2003; Loh et al. 2006). It contains a homeodomain and two intrinsically disordered regions (IDRs) at the N- and C-terminus (Figure 4A,B). The Nanog homeodomain is highly conserved and mediates DNA binding (Schuff et al. 2012; Theunissen et al. 2011; Jauch et al. 2008). The *in vivo* function of the disordered N- and C-terminus, however, is less clear. Here, to investigate which domains of Nanog play a role in the formation of the two *mir430* transcription bodies, we generated a series of mutant Nanog constructs. We deleted the homeodomain (ΔHD), both IDRs at the same time (HD only) and each of the IDRs separately (ΔN and ΔC) (Figure 4C). Because the lack of *mir430* transcription bodies in the *Nanog* mutant can be rescued by injecting RNA encoding full length Nanog into 1-cell stage embryos (Figure 4D), injecting RNA derived from these mutant constructs allowed us to probe the importance of the homeodomain and the disordered regions in the formation of transcription bodies. We observed that in absence of the homeodomain (ΔHD) no transcription bodies formed. This suggests that the DNA binding domain of Nanog is required for transcription body formation. It is, however, also possible that the Nanog homeodomain contributes to the formation of transcription bodies in other ways. To make sure that the effect of losing the homeodomain is the consequence of affecting DNA binding, we also generated a series of single-point mutations to alanine in conserved residues of the HD that have been previously shown to abrogate DNA binding (R247A, T241A, N245A; Figure 4D; (Hayashi et al. 2015)). Injecting RNA encoding the R247A mutant did not rescue the formation of the *mir430* transcription bodies (Figure 4D), confirming that DNA binding is required for transcription body formation. We note that two other point-mutations that also are known to affect DNA binding (HD T241A and N245A), showed a delay in transcription body formation, but they did form eventually (Figure 4D). We hypothesize that these mutants retain some DNA binding capacity. Next, to test whether DNA binding is sufficient, we injected just the homeodomain in *Nanog* mutant embryos (Figure 4C, HD only). Here, we observed no transcription bodies (Figure 4D). Thus, while DNA binding is required, it is not sufficient for the formation of transcription bodies. To test the importance of N- and C-terminus, we deleted them independently (Figure 4C; ΔN and ΔC, respectively). In both cases, we observed transcription bodies (Figure 4D). The fraction of embryos in which we see rescue of transcription body formation at 128-cell suggests that N-terminus is more important than the C-terminus, which may be related to the role of the N-terminus in Nanog dimerization (Schuff et al. 2012). In addition to the absence of the two prominent *mir430* transcription bodies, *Nanog* mutants also display gastrulation defects (Figure 4E; Veil et al. 2018; Gagnon et al. 2018; Pálfy et al. 2020). This is a consequence of the important role of Nanog in developmental gene expression (Lee, Bonneau, and Giraldez 2014; Gagnon, Obbad, and Schier 2018; Veil et al. 2018). Similar to transcription body formation, these developmental defects can be rescued by injection of full-length Nanog RNA at the 1-cell stage. Interestingly, we observed that the mutant forms of Nanog that were able to rescue transcription body formation, were also able rescue the gastrulation phenotype (Figure 4E). Because Nanog plays an important role in developmental gene expression beyond the *mir430* locus, this result suggests that the ability of Nanog to organize transcription bodies may be generally important for Nanog function. From the work presented here, we can conclude that the Nanog homeodomain, and either one of its IDRs, are required to recruit RNA Pol II and form transcription bodies.

**Figure 4.**
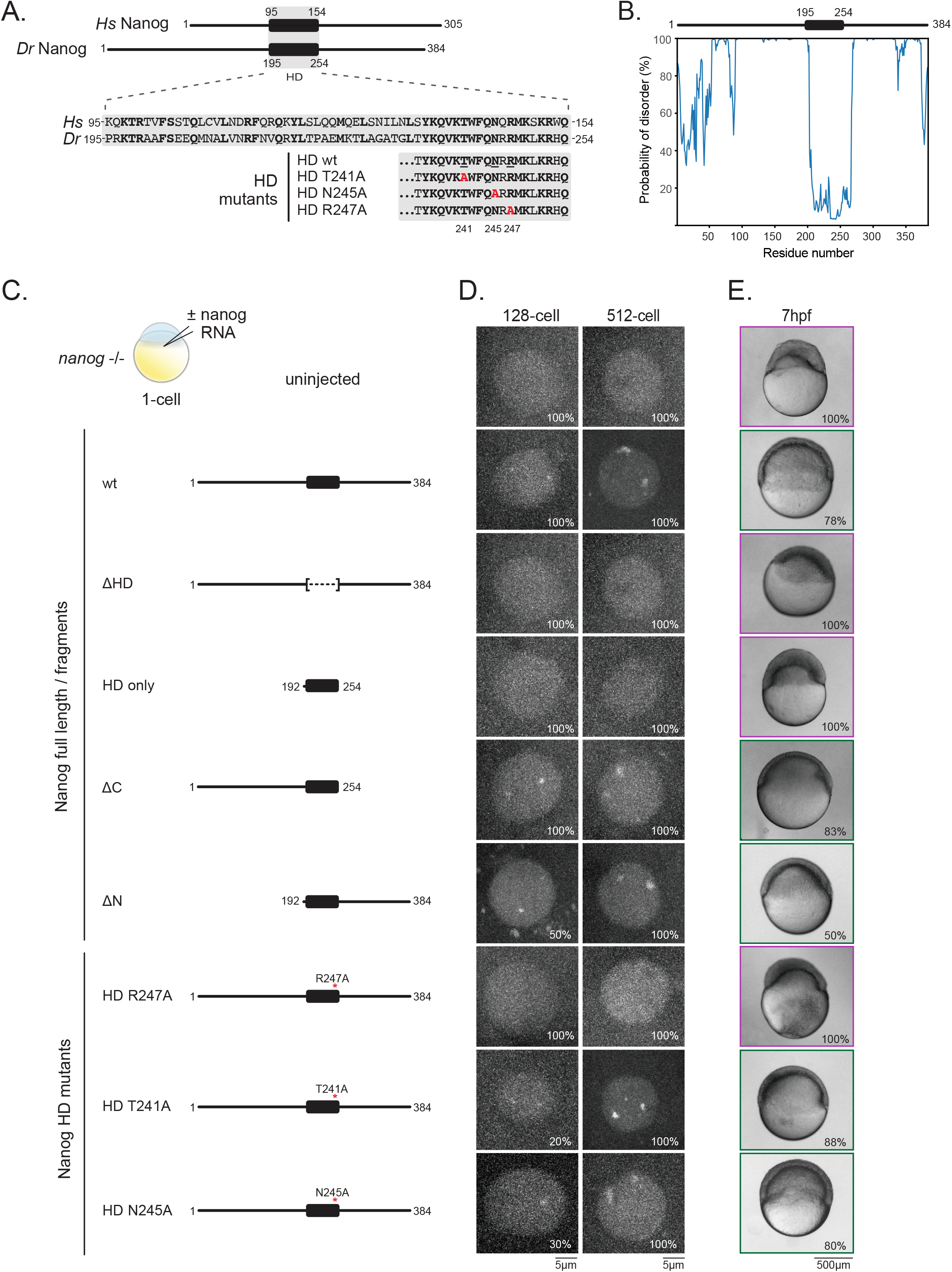
Nanog DBD as well as IDR are required to seed transcription bodies. **A**. Domain structure of human (Hs) and zebrafish (Dr) Nanog. Conserved residues in the homeodomain are indicated in bold, homeodomain mutants used are indicated. HD, homeodomain. **B**. Order and disorder in zebrafish Nanog protein as predicted by ODiNPred (Dass, Mulder, and Nielsen 2020). **C**. Schematic representation of experimental set-up and constructs used. **D**. Visualization of *mir430* transcription bodies with Ser2P Fabs at 128- and 512-cell in embryos injected with construct indicated to the left. **E**. Visualization of developmental stage reached 7 hours post fertilization (hpf) upon injection of construct indicated on the left. Purple and green boxes indicate arrested and rescued phenotype, respectively. In D and E, percentages indicate how often the shown phenotype is observed. For D and E, N ≥ 3 and n ≥ 18.

In this study we analyzed the assembly of two transcription bodies that mark the beginning of zygotic transcription in the zebrafish embryo. In these *mir430*-seeded transcription bodies, the two pluripotency transcription factors Nanog and Sox19b, as well as the initiating and elongating form of RNA polymerase II cluster. In a highly stereotyped process, Nanog clusters first, and this is required to bring in Sox19b and activate transcription. The organizing role of Nanog requires its DNA binding domain as well as either one of its IDRs. Once transcription is well underway and RNA accumulates, transcription factor clusters disassemble.

Our observation that Nanog and Sox19b form clusters in the nucleus is in agreement with previous observations of transcription factor clustering (Tsai et al. 2017; Mir et al. 2017; Nair et al. 2019; Sabari et al. 2018; Li et al. 2020; Ma et al. 2021). While transcription factors have been shown to be able to recruit Mediator (Boija et al. 2018) and even RNA Pol II in artificial systems (Chong et al. 2018), it has been unclear how transcription bodies form *in vivo*. Here we show that transcription factors can organize transcription bodies.

A role for transcription factors in scaffolding transcription bodies has previously been proposed (Sabari et al. 2018; Boija et al. 2018). In this model, transcription factors bind to DNA with their DNA binding domain, while recruiting transcriptional co-activators with their IDRs. Our results provide experimental support for this model. Both the Nanog DNA-binding domain as well as one if its IDRs is required to scaffold transcription bodies. Interestingly, we found that Nanog does not only recruit RNA Pol II, but also Sox19b. The coincidence of Sox19b and Nanog, however, is not observed in all Nanog clusters, suggesting a high level of specificity in the formation of transcription bodies. Future work will clarify how such specificity is achieved.

## ACKNOWLEDGEMENTS

We thank the members of the Vastenhouw and Brugues labs as well as the MZT community for helpful feedback and stimulating discussions, Shivali Dongre and Noémie Chabot for comments on the manuscript, and the following facilities and services of the MPI-CBG for their support: fish facility, light microscopy facility, computer department, scientific computing. This work was supported by the Max Planck Society, the University of Lausanne, the Volkswagen foundation ‘Life’ initiative (94773 to NLV), the DFG SPP ‘Molecular mechanisms of functional phase separation’ (VA 1209/2-1 to KK, NLV), and the Japan Society for the Promotion of Science KAKENHI (JP18H05527 and JP21H04764 to HK, and JP20K06484 to YS).

## STAR METHODS

### Zebrafish and molecular biology approaches

#### Zebrafish husbandry and manipulation

Zebrafish were maintained and raised under standard conditions. Wild-type (TLAB) and mutant (*Nanog*-/- (Veil et al. 2018) and *mir430*-/- (Figure S1)) embryos were dechorionated with Pronase (Pronase E, 107433, Merck, Sigma Aldrich) immediately upon fertilization, synchronized and allowed to develop to the desired stage at 28°C. Stage was determined by morphology and corroborated by cell counting and measuring distances between nuclei. A-amanitin (A2263; Sigma) was injected at the 1-cell stage at a concentration of 0.2 ng per embryo. Fabs were injected at the 1-cell stage at concentrations indicated in Table 1. Bright-field images of whole embryos were acquired on a stereomicroscope (Olympus, SZX-12) equipped with an appropriate CCD camera (Qimaging, 12bit Cooled Mono).

**Table 1.**
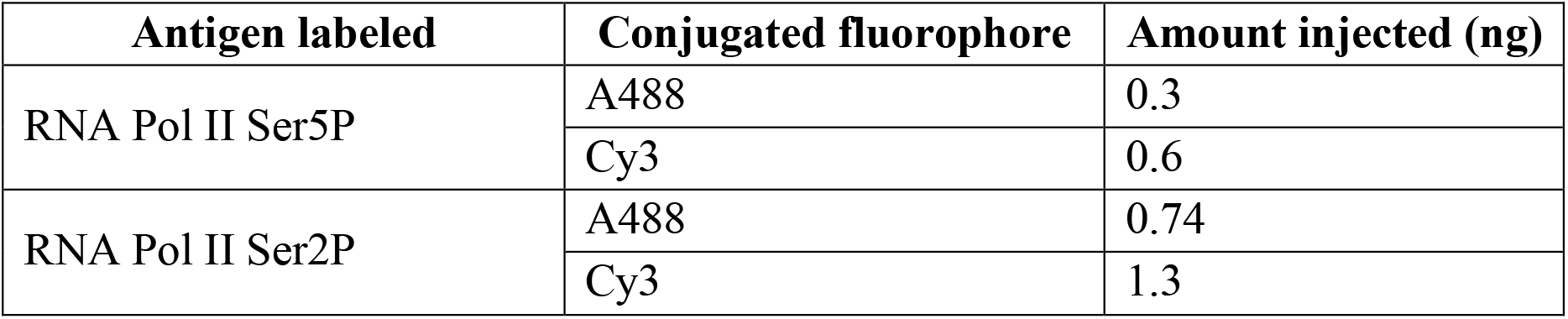
Fluorescently labeled antibody fragments (Fabs) used.

#### mRNA production and injection

mRNA was synthesized using the Ambion mMESSAGE mMACHINE SP6 Transcription Kit (AM1430; ThermoFisher Scientific, Waltham, MA). cDNAs were cloned into a pCS2+ vector with a C-terminal tag (see Table 2 for a complete list). mRNA was injected into the cell at the 1-cell stage in amounts indicated in Table 2. Please note that for the rescue experiments, the amount of RNA injected was based on the 120pg that was published to be required to rescue *Nanog* mutant embryos (Veil et al. 2018; Pálfy et al. 2020) and corrected for the differently sized constructs.

**Table 2.** mRNA constructs used.

| cDNA | C-term tag | Experiment type | Amount injected (pg) |
| --- | --- | --- | --- |
| Nanog | mNeonGreen | Visualization of Nanog | 100 |
| Sox19b | mNeonGreen | Visualization of Sox19B | 150 |
| Pou5f3 | mNeonGreen | Visualization of Pou5f3 | 150 |
| Nanog | HA | Rescue of *Nanog*<sup>-/-</sup> mutants (line propagation) | 120 |
| Nanog | HA | Rescue of transcriptional bodies and/or phenotype in *Nanog*<sup>-/-</sup> embryos | 120 |
| HDonly | HA |  | 50 |
| ΔN | HA |  | 70 |
| ΔC | HA |  | 70 |
| HD T241A | HA |  | 120 |
| HD N245A | HA |  | 120 |
| HD R247A | HA |  | 120 |

#### Generation of mir430 mutants

Two-part gRNAs consisting of crRNAs and tracrRNA were used to target up- and downstream of the *mir430* genomic locus. crRNAs were designed using CHOPCHOP ((Montague et al. 2014); Table 3). Both crRNAs and tracrRNA (IDT, Alt-R® CRISPR-Cas9 tracrRNA) were ordered from Integrated DNA technologies (IDT), reconstituted in 100μM nuclease-free water and hybridized at 3μM concentration by incubating them at 95°C for 5 minutes. To prepare the injection solution, Alt-R S.p. Cas9 Nuclease (IDT) was diluted in Cas9 working buffer (20mM HEPES, 150mM KCl, pH 7.5) to a final concentration of 1.35μg/μL, mixed with the two hybridized gRNAs in a 2:1:1 ratio, incubated at 37°C for 10 minutes and brought to room temperature. 1nL was injected into wild type embryos at the 1-cell stage.

**Table 3.**
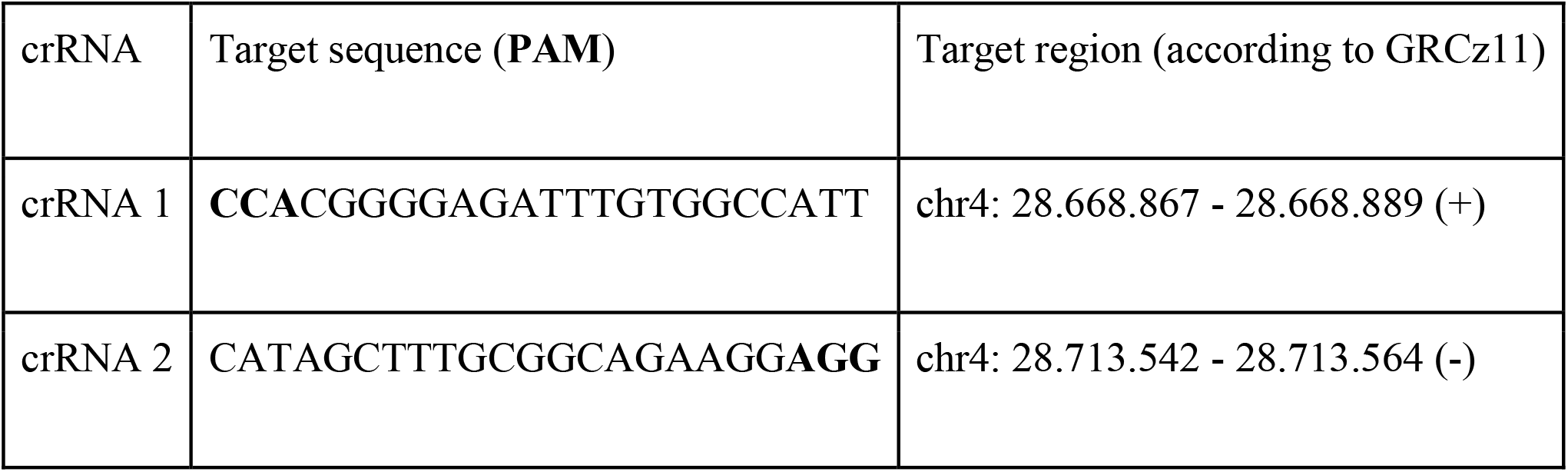
crRNAs used to generate *mir430* mutant fish.

Founder fish were identified by incrossing potential founders and screening the progeny for the presence of the mutant band (Figure S1). Heterozygous fish were obtained from founder fish by outcrossing them with wild type fish. Homozygous fish were then obtained by incrossing either heterozygous or homozygous fish and injecting the obtained embryos with 2nL of rescue solution (10μM of miR430a, miR430b and miR430c duplex (Giraldez 2006) in 1x siRNA solution (300mM KCl, 30mM HEPES, 1mM MgCl2, brought to pH 7.5 with KOH), see (Giraldez 2006). To assess the genotype, adult fish were genotyped by anesthetizing them with MESAB solution (168mg/L Tricaine (Sigma-Aldrich, #A5040) and 168-336mg/L NaHCO_3_) and quickly clipping a piece of their fin with a scalpel. Clipped fins were collected in PCR plates and PCR-grade genomic DNA was then obtained through the Hot Shot method ((Truett et al. 2000): In brief, fins were incubated for 45 minutes at 95°C in 100μL of 50mM NaOH. After brief cooling at 4°C, 10μL of 1M Tris-HCl pH 8.0 was added, and 1μL of the obtained lysate was used to set up a 20μL genotyping PCR reaction mix (see Table 4 for primer sequences and running conditions). In-house made Taq (MPI-CBG) and NEB Taq Polymerase (NEB, #M0273) were used. PCR reactions were run on agarose gels alongside the 1kb Plus DNA Ladder (NEB, #N3200), see Figure S1.

**Table 4.**
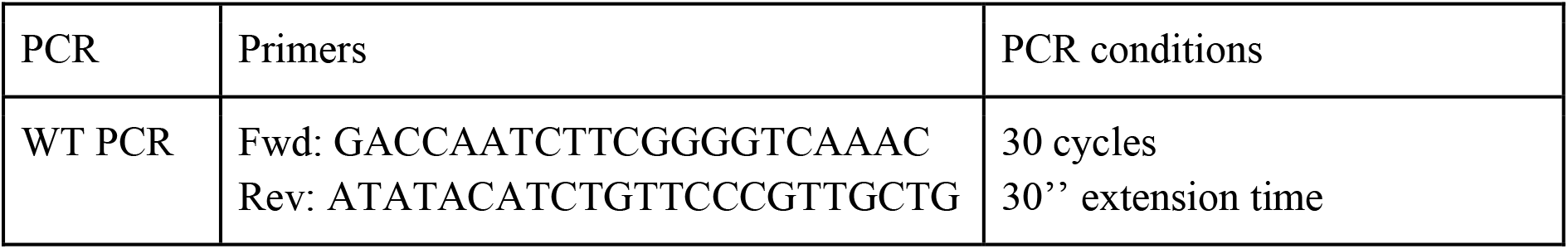
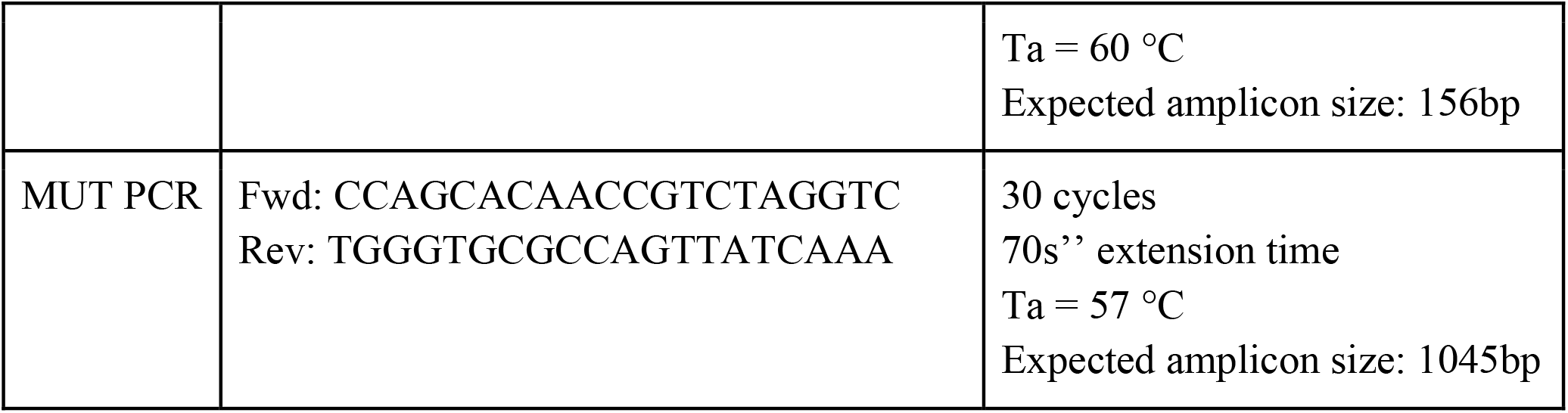
Primers used to genotype *mir430* mutant fish.

### Live microscopy

#### Preparation of fragmented antibodies (Fabs) for use in live-cell microscopy

Fluorescently labeled fragmented antibodies (Fabs) specific to Pol II Ser5P and Pol II Ser5P were prepared from monoclonal antibodies specific to Pol II Ser5 and Ser2 phosphorylation (Hayashi-Takanaka et al. 2011; Stasevich et al. 2014; Kimura and Yamagata 2015). Monoclonal antibodies were digested with Ficin or Papain (ThermoFisher Scientific), and Fabs were purified through protein A-Sepharose columns (GE Healthcare) to remove Fc and undigested IgG. After passing through desalting columns (PD MiniTrap G25; GE Healthcare) to substitute the buffer with PBS, Fabs were concentrated up to >1 mg/ml using 10 k cut-off filters (Amicon Ultra-0.5 10 k; Merck), Fabs were conjugated with Alexa Fluor 488 (Sulfodichlorophenol Ester; ThermoFisher Scientific) or Cy3 (N-hydroxysuccinimide ester monoreactive dye; GE Healthcare) to yield ∼1:1 dye:protein ratio. After the buffer substitution with PBS, the concentration was adjusted to ∼1 mg/ml.

#### Preparation of embryos for use in live-cell microscopy

At 16- to 32-cell stage, embryos were mounted for live imaging in 0.7% UltraPure low-melting-point (LMP) Agarose (ThermoFisher 16520050) dissolved in Danieau’s supplemented with iodixanol (OptiPrep, STEMCELL Technologies 07820) to match a refractive index of 1.3615 (Boothe et al. 2017). Ibidi glass-bottom dishes (μ-Dish 35 mm, high Glass Bottom, 81158) were used. Embryos were brought closer to the coverslip surface by keeping the dish upside down until agarose solidified. Then, the embryos were coated with agarose to avoid drying of the sample. Finally, embryos were brought to the microscopy room in a light-protected transfer box. Imaging commenced from around 32-cell stage.

#### Live imaging by spinning-disk microscopy

All live imaging experiments (except for the data presented in Figure 1H) were performed using the spinning disk Andor Revolution platform with Borealis extension, equipped with an Olympus silicone oil-immersion objective (UPlanSApo 60x, NA 1.30, OLYMPUS), recording with simultaneous imaging of two fluorophores with the help of two iXon Ultra 888 EMCCD cameras. Data for Figure 1H was obtained using a Nikon eclipse Ti2 with a Yokogava CSU-W1 Spinning Disk unit at 28°C (okolab temperature control and microscope enclosure) equipped with a Nikon 60x/1.2 Plan Apochromat VC water objective. Images were captured in parallel on two Photometrics Prime 95B cameras. For both set-ups, time-lapses were recorded over periods of up to 160 min, during which the embryos divided, suggesting no obvious phototoxicity.

### Image processing and analysis

#### Software used for image handling and analysis

Microscopy image handling was done using Fiji (Schindelin et al. 2012) and Jupyter Notebooks running custom Python code. Further data processing was carried out with TrackMate (Tinevez et al. 2017), Imaris (v 9.9.0), and R Studio. The resulting figures were prepared using ScientiFig (Aigouy and Mirouse 2013) and movie annotation plugin (Daetwyler, Modes, and Fiolka 2020).

#### Image pre-processing with Noise2Void

In preparation for tracking the nuclear clusters, images were denoised with Noise2Void (Krull, Buchholz, and Jug 2019). The network was trained on and applied to the raw spinning disk confocal data in full 3D with both color channels being present. After denoising, the resulting volumes are automatically max-projected and the 3D and max-projected 2D data stored for further downstream processing. The used notebooks are adapted from Krull, Buchholz, and Jug 2019 and can be shared with interested parties upon reasonable or unreasonable requests.

#### Signal normalization

The denoised and max-projected 2D image data was normalized with the help of a custom-made Python script using the napari image-viewing (Sofroniew et la. 2022) for visualization and interactive cropping. The normalization was implemented using Contrast limited adaptive histogram equalization, CLAHE (Zuiderveld 1994) and also corrected for nuclear drift introduced by cell motion and division. Normalized and motion corrected 2D data was saved for further downstream processing. The script can be shared with interested parties upon reasonable or unreasonable requests.

#### Tracking with TrackMate

Nuclei were segmented in the denoised and max-projected image data using Otsu thresholding. Connected components in the thresholded images define the regions of interest (ROI) we subsequently saved for cluster detection and tracking. Cluster identification was carried out using the LoG (Laplacian of Gaussian) detector of TrackMate (Tinevez et al. 2017). The estimated object diameter was set to 1μm for TFs, and 1μm (64-cell) or 2μm (for 128- and 256-cell stage) for RNA polymerase clusters. Following automatic detection, identified spots were tracked using TrackMate’s LAP (Linear Assignment Problem) tracker. Tracking of transcription bodies (RNA Pol II Ser5P and Ser2P) was performed semi-automatically to further improve the quality of the obtained tracks. All detection and tracking results were manually checked to avoid downstream analysis of faulty tracks. Remaining tracking errors have been curated manually. The tracking of TF clusters and the colocalization with transcription body was also performed manually.

#### Determining developmental stage and mid-point between interphases

Developmental stages (64, 128, 256 and 512 cell stages) were determined based on sample preparation time and the distances between nuclei. This method is very reliable as the inter-nuclear distances in these early stages are highly stereotypic. The precise point in time at which we separate consecutive cell stages was visually defined at mitosis (anaphase).

#### Measuring cluster size with Imaris

The area of RNA pol II Ser5P clusters in 2D max-projected images of the 64- to 256-cell stage was measured using Imaris. We used the built-in “Spots” model with the diameter parameter set to 1.9μm. The possibility of variable sizes between different timepoints in the timelapse was enabled via the “local contrast values” parameter. Spots were filtered according to the “median intensity” parameter of the Pol II Ser2P channel (set to 5e4, and manually adjusted if required). The “region growth threshold” parameter was typically set to 80.7 (and also manually adjusted if needed). Manually set parameters were based on visual examination, such that the full area of the focus was included in subsequent measurements. Finally, the diameter and area measurements were determined via the unsuitably called “volume” parameter. In order to track the changes of the same clusters throughout the entire timelapse, identified Spots were tracked using a maximum inter-timepoint distance of 3.49μm. We typically report transcription body areas at the mid-point of each cell-stage interphase. When the growth within one interphase was plotted all detected spot-sizes were used.

### Data analysis

#### Sample size determination

A minimum of 3 biological and 3 technical replicates was generated for each experiment. The number of experimental replicates (N) as well as the number of measured nuclei (n) are reported for each conducted experiment individually in the respective figure legend.

#### Statistics and plotting with RStudio software

For tracking, each of the channels was treated as a separate entity with Trackmate. Therefore, tracks were plotted and connected via XY location in the image. For the main analysis and table modifications, the following R packages were used: “ggplot2”, “ggpubr”, “tidyverse”, “dplyr” and “rstatix” (Kassambara 2020; Kassambara 2020; Wickham 2016, 2017; Wickham et al. 2015). P-values were calculated using the paired non-parametric Wilcoxon test (Wilcoxon 1945; Mann and Whitney 1947). All scripts can be shared with interested parties upon reasonable or unreasonable requests.

## SUPPLEMENTAL FIGURE LEGENDS

**Figure S1.**
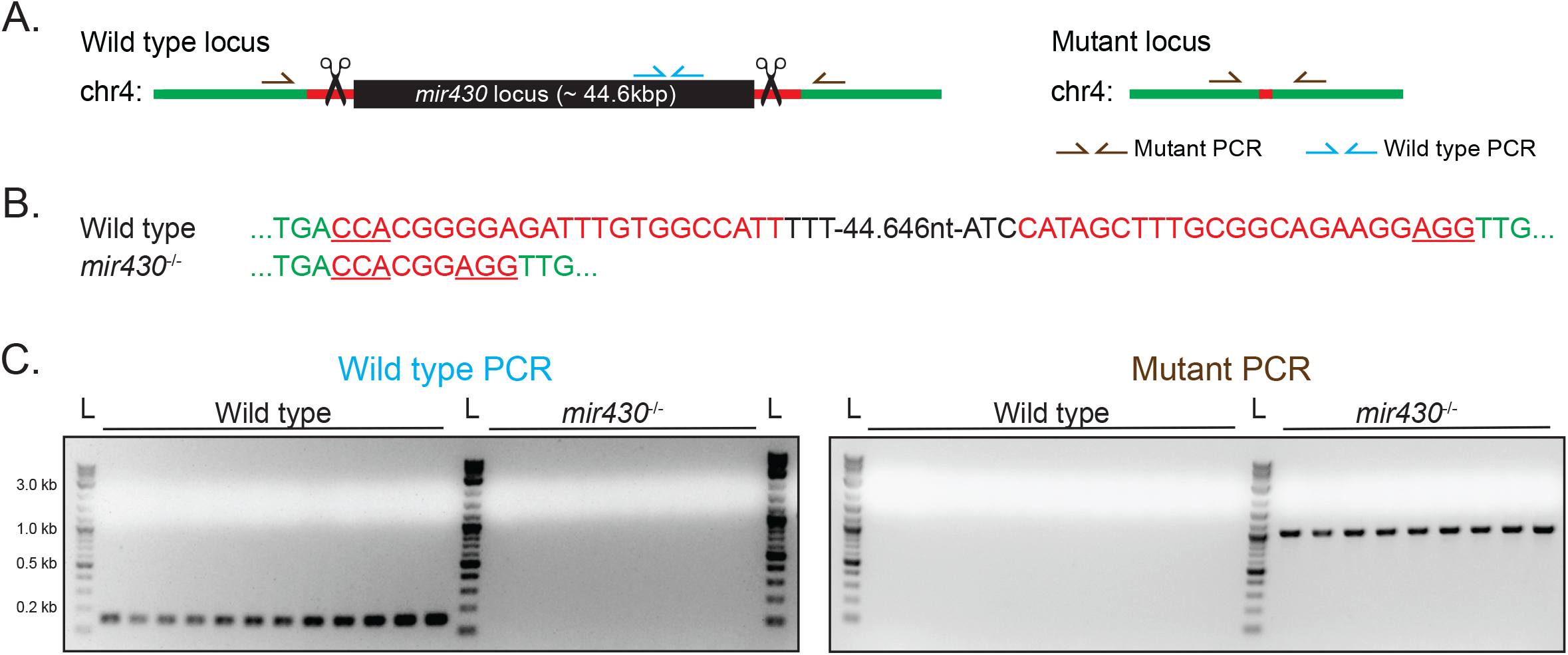
The generation of *mir430* mutant fish. **A**. Schematic representation of the *mir430* locus, the gRNA target sites (red) and the two primer pairs used for genotyping (brown and azure). **B**. Sequence of the mutant allele as determined by Sanger sequencing of the PCR band obtained by the Mutant PCR. gRNA target sites are shown in red, and the PAM sites are underlined. **C**. Representative agarose gel showing the specificity of the Wild type and Mutant PCRs. Wild type samples generate only the wild type PCR band, while homozygous deletion samples generate only the mutant PCR band. L: DNA Ladder (NEB 1kb Plus).

**Movie S1. Pol II Ser5P and Pol II Ser2P clusters at 64-cell stage**. Spinning disk confocal microscopy timelapse of the 64-cell stage nucleus shown in Figure 1B. Cyan arrowheads indicate Pol II Ser5P and Ser2P clusters that coincide. Magenta arrowheads indicate Pol II Ser5P clusters that do not acquire Pol II Ser2P signal. Scale bar is 5μm. The movie represents maximum intensity projections (MIPs) through the Z-stack, time indicated in minutes.

**Movie S2. Pol II Ser5P and Pol II Ser2P clusters at 128-cell stage**. Spinning disk confocal microscopy timelapse of the 128-cell stage nucleus shown in Figure 1B. Cyan arrowheads indicate Pol II Ser5P and Ser2P clusters that coincide. Magenta arrowheads indicate Pol II Ser5P clusters that do not acquire Pol II Ser2P signal. Scale bar is 5μm. The movie represents maximum intensity projections (MIPs) through the Z-stack, time indicated in minutes.

**Movie S3. Pol II Ser5P and Pol II Ser2P clusters at 256-cell stage**. Spinning disk confocal microscopy timelapse of the 256-cell stage nucleus shown in Figure 1B. Cyan arrowheads indicate Pol II Ser5P and Ser2P clusters that coincide. Magenta arrowheads indicate Pol II Ser5P clusters that do not acquire Pol II Ser2P signal. Scale bar is 5μm. The movie represents maximum intensity projections (MIPs) through the Z-stack, time indicated in minutes.

**Movie S4. Nanog and Pol II Ser5P clusters at 128-cell stage**. Spinning disk confocal microscopy timelapse of the 128-cell stage nucleus shown in Figure 2A. Left (Nanog): Cyan arrowheads indicate Nanog clusters that will acquire Pol II Ser5P, magenta arrowheads indicate Nanog clusters that do not. Right (Pol II Ser5P): Cyan arrowheads indicate the two *mir430* transcription bodies, magenta arrowheads indicate Pol II Ser5P clusters that do not acquire elongation signal. Scale bar is 5μm. The movie represents maximum intensity projections (MIPs) through the Z-stack, time indicated in minutes.

**Movie S5. Sox19b and Pol II Ser5P clusters at 128-cell stage**. Spinning disk confocal microscopy timelapse of the 128-cell stage nucleus shown in Figure 2A. Left (Sox19b): Cyan arrowheads indicate Sox19b clusters that will acquire Pol II Ser5P. Right (Pol II Ser5P): Cyan arrowheads indicate the two *mir430* transcription bodies. Scale bar is 5μm. The movie represents maximum intensity projections (MIPs) through the Z-stack, time indicated in minutes.

**Movie S6. Pou5f3 and Pol II Ser5P clusters at 128-cell stage**. Spinning disk confocal microscopy timelapse of the 128-cell stage nucleus shown in Figure 2A. Left (Pou5f3): Magenta arrowheads indicate Pou5f3 clusters. Right (Pol II Ser5P): Cyan arrowheads indicate the two *mir430* transcription bodies, magenta arrowheads indicate Pol II Ser5P clusters that do not acquire elongation signal. Scale bar is 5μm. The movie represents maximum intensity projections (MIPs) through the Z-stack, time indicated in minutes.

**Movie S7. Nanog and Pol II Ser5P clusters at 128-cell stage in α-amanitin treated embryos**. Spinning disk confocal microscopy timelapse of the 128-cell stage nucleus shown in Figure 2C. Left (Nanog): Cyan arrowheads indicate Nanog clusters that acquire Pol II Ser5P, magenta arrowheads indicate some of the Nanog clusters that do not. Right (Pol II Ser5P): Cyan arrowheads indicate the two transcription bodies. Scale bar is 5μm. The movie represents maximum intensity projections (MIPs) through the Z-stack.

**Movie S8: Sox19b and Pol II Ser5P clusters at 128-cell stage in α-amanitin treated embryos**. Spinning disk confocal microscopy timelapse of the 128-cell stage nucleus shown in Figure 2C. Left (Sox19b): Cyan arrowheads indicate Sox19b clusters that acquire Pol II Ser5P. Right (Pol II Ser5P): Cyan arrowheads indicate two transcription bodies. Scale bar is 5μm. The movie represents maximum intensity projections (MIPs) through the Z-stack.

**Movie S9: Sox19b clusters at 128-cell stage in WT embryo**. Spinning disk confocal microscopy timelapse of nucleus at 128-cell stage shown in Figure 3C (Sox19b in WT). Cyan arrowheads indicate two Sox19B clusters. Scale bar is 5μm. The movie represents maximum intensity projections (MIPs) through the Z-stack.

**Movie S10: Absence of Sox19b clusters at 128-cell in *nanog-/-* embryo**. Spinning disk confocal microscopy timelapse of nucleus at 128-cell stage shown in Figure 3C (Sox19b *nanog-/-*). No Sox19b clusters can be observed. Scale bar is 5μm. The movie represents maximum intensity projections (MIPs) through the Z-stack.

**Movie S11: Pol II Ser5P clusters at 128-cell in WT embryo**. Spinning disk confocal microscopy timelapse of nucleus at 128-cell stage shown in Figure 3D (Pol II Ser5P in WT). Cyan arrowheads indicate the two *mir430* transcription bodies. Scale bar is 5μm. The movie represents maximum intensity projections (MIPs) through the Z-stack.

**Movie S12: Absence of Pol II Ser5P clusters at 128-cell in *nanog-/-* embryo**. Spinning disk confocal microscopy timelapse of nucleus at 128-cell stage shown in Figure 3C (Pol II Ser5P in *nanog-/-*). No Ser5P clusters can be observed. Scale bar is 5μm. The movie represents maximum intensity projections (MIPs) through the Z-stack.

